# Delayed Center of Mass Feedback in Elderly Humans Leads to Greater Muscle Co-Contraction and Altered Balance Strategy under Perturbed Balance: a Predictive Musculoskeletal Simulation Study

**DOI:** 10.1101/2023.12.18.572216

**Authors:** Rachel Jones, Neethan Ratnakumar, Kübra Akbaş, Xianlian Zhou

**Affiliations:** Department of Biomedical Engineering, New Jersey Institute of Technology, Newark, NJ, USA

## Abstract

Falls are one of the leading causes of non-disease death and injury in the elderly, often due to delayed sensory neural feedback essential for balance. This delay, challenging to measure or manipulate in human studies, necessitates exploration through neuromusculoskeletal modeling to reveal its intricate effects on balance. In this study, we developed a novel three-way muscle feedback control approach, including muscle length feedback, muscle force feedback, and enter of mass feedback, for balancing and investigated specifically the effects of center of mass feedback delay on elderly people’s balance strategies. We conducted simulations of cyclic perturbed balance at different magnitudes ranging from 0 to 80 mm and with three center of mass feedback delays (100, 150 & 200 ms). The results reveal two key points: 1) Longer center of mass feedback delays resulted in increased muscle activations and co-contraction, 2) Prolonged center of mass feedback delays led to noticeable shifts in balance strategies during perturbed standing. Under low-amplitude perturbations, the ankle strategy was predominantly used, while higher amplitude disturbances saw more frequent employment of hip and knee strategies. Additionally, prolonged center of mass delays altered balance strategies across different phases of perturbation, with a noticeable increase in overall ankle strategy usage. These findings underline the adverse effects of prolonged feedback delays on an individual’s stability, necessitating greater muscle co-contraction and balance strategy adjustment to maintain balance under perturbation. Our findings advocate for the development of training programs tailored to enhance balance reactions and mitigate muscle feedback delays within clinical or rehabilitation settings for fall prevention in elderly people.

## Introduction

According to the Center for Disease Control and Prevention (CDC), falls are the leading cause of non-disease death in individuals over the age of 65 [1, 2]. Even if a fall does not result in death, it can cause serious injury – up to 20% of falls result in bone fractures or head injuries [3–5] – and put the individual at increased risk: after an initial fall, the likelihood of experiencing another fall is doubled [6]. Maintenance of standing balance is a necessary component to quality of life. Not being able to stand steadily for extended periods of time minimizes the daily activities one can perform [7]. It is well established that elderly people have delayed sensory motor responses compared to the younger population [8, 9], this change in reaction time diminishes balance capacity and puts elderly people at an increased risk for falls [10]. To protect elderly people – a vulnerable population group – a proper understanding of the balance strategies adapted under the decline of sensory motor capabilities is needed.

Human balance is not a function of conscience control, rather it involves a complex interplay of various reflexes of the central nervous system (CNS), including the vestibular, proprioceptive, and visual systems [11]. These systems provide sensory information about the position and movement of the body to the brain and help to involve various reflexes and controls to maintain balance. Proprioception is important for postural control because it has a lower threshold for response than vestibular or visual systems [12]. Muscle spindles – a part of the proprioceptive system – are sensory receptors that detect changes in muscle lengths and the speed at which these changes occur, muscle stretch will cause the spindles to activate the stretch reflex and signal the muscle to contract to prevent damage to muscle fibers and maintain balance. These proprioceptive feedback pathways are often damaged in elderly people through age-related diseases such as diabetes and arthritis or morphological changes to the sensory fibers [12, 13]. Elderly people have been found to have slower voluntary muscle contraction due to slower neuron signaling and reaction times in comparison to the younger population [10, 14–16]. Additionally, elderly people tend to rely more heavily on visual feedback, which is less sensitive than proprioception and can lead to difficulties with balance control [17, 18]. As human bodies age, there is a decline in sensory systems feedback [19–21], but timely response to sensory feedback is paramount to balance control [22].

For proper balance maintenance, the body must keep the Center of Mass (COM) within the Base of Support (BoS), which is the area of contact between the feet and the ground. When the COM is outside the BoS, the body will begin to fall, and the sensory systems in the body will initiate feedback mechanisms to restore balance. If the COM moves too far outside of the BoS, balance will not be regained without taking a step – possibly leading to a fall. The area that balance can be recovered from varies based on a number of factors (i.e., subject paying attention, expectation of perturbation, prior experience, sensory response time, etc.) that differ from person to person [23]. Another important indicator of balance is the movement of the Center of Pressure (COP), which is a point within the BoS where the summed ground contact forces acting on the feet are applied. The movement of the CoP is directly influenced by the COM movement [24]. In the study of human balance reduced order models, such as the inverted pendulum (IP), serve as useful tools for understanding whole-body kinematics and kinetics [24–27]. IP-based models have been widely used to study sagittal plane kinematics/kinetics for balance and gait in bipeds, as well as in designing controllers, making them a popular initial approach. However, these models do not incorporate muscles and neuromuscular control into the picture.

While balance during gait can be maintained by lateral shifting foot placement – thereby widening the BoS – standing balance has a static BoS and relies on muscle activations to bring the COM back into BoS [28–30]. Which muscles are activated and by how much depends on the movement of the COM. Large deviations from the center of the BoS cause activations in the muscles surrounding the hip, whereas smaller deviations are corrected via shank muscles controlling the ankle angle [25, 26, 31–33]. During perturbed balance maintenance, when the body is suddenly moved or the BoS is shifted, the muscles respond rapidly to maintain balance and prevent falls. However, there is a sensory feedback delay in the response due to the time it takes for the sensory information to be processed and the motor commands to be sent to the muscles. Typical muscle activation delays due to feedback differ by muscle, but are in the range of 35-100 ms for lower limb muscles [34]. Sensory feedback delay is an important consideration in understanding balance control and fall prevention in the vulnerable elderly population.

Previous balance studies typically focused on quantifying the human reaction (e.g., body movement or electromyography (EMG)) to perturbation. Van der Kooij and Vlugt measured COM and ankle torque of human subjects on a perturbed platform and found that the sensory feedback loop depends on estimations of body position [35]. In a study on perturbed walking by Olivera *et al.* [36], the authors used EMG signals to define motor modules, or synergies, for gait, and then noted the changes when the subjects adapted to unexpected perturbations during gait. While the motor synergies were largely unaffected by perturbation, the activations of those muscles were significantly different in perturbation trials. Although these studies provide significant insight regarding balance kinematics and muscle activity, they cannot study the effects of biological variables such as sensory feedback delay on balance in human subjects. Therefore, computational musculoskeletal (MSK) modeling has emerged as a prominent tool for investigating the neuromotor elements of human balance. In a study by Aftab *et al.* [25], the authors developed a predictive computational modeling framework to detect balance recovery strategies based on COP and upper body inertia. In pursuit of physiological mimicry, Aoi et al. created a complex spinal reflex controller that closely mirrors the pathways and methods used by the human body to control balance [37]. The development of this complex controller brings up the question of physiological accuracy vs computational ease. A close approximation of physiological neural networks results in an excessive level of complexity, making it unfeasible to explore different scenarios with the computational resources at hand.

Previous studies have used both discrete or impulse perturbance and continuous perturbation to evaluate a subject’s ability to react and other aspects of perturbed balance recovery. Some researchers have investigated the adaptation of balance strategies during continuous perturbation. In a study on balance strategies by Akram *et al*. [38], the authors subjected participants to continuous rotational perturbances during quiet standing, finding multi-joint balance strategies are necessary for balance during rotational perturbations. In a study on vision-based balance by Buchanan and Horak [32], continuous sinusoidal perturbations were applied at a fixed magnitude with variations in frequency to quantify the relationship between translational frequency and postural patterns. Five minutes of continuous perturbations via waist pulls was used in a study by Camernik *et al.* [39], which evaluated the effects of handles on posture, finding lower activations in the major posture stabilizing muscles when a handle was used. While these studies have contributed to the scientific knowledge regarding balance, none explored sensory feedback in their analysis of balance strategies. Studying balance strategies during continuous perturbation, while being able to adjust the COM feedback delay can provide new insights about aging and balance.

To keep the body upright, the MSK system and CNS are constantly processing sensory feedback and sending out corrections. These feedback systems need to be included in the framework to design a reliable human balance model. In this study, we intended to explore how the length of sensory feedback delay affects balance strategies. To achieve this objective, we used a MSK optimization approach to simulate the effects of COM feedback control delay in the response from the MSK system during perturbation. We hypothesized that longer COM feedback delay would result in greater activations and co-contraction in the major stabilizing muscles (signaling age induced joint instability) and alteration of balance strategies. The overall motivation of this study was to use a MSK model with controller optimization to explore the effects of COM feedback delay on balance control, which sheds light on the balance control changes implemented by the human body to adjust for longer sensory feedback delays, like those seen in the elderly population.

In this paper, we will first introduce a novel three-way controller approach used for muscle reflexes during balancing and explain how the controllers reflect the neurophysiological control process, followed by a brief introduction of the method to solve the optimization problem of muscle control. Then we will present the results of joint angle, muscle activation and co-contraction, relative COM and CoP movement, joint moment, and balance strategy under 10 perturbation conditions and three COM delay conditions. Finally, we will discuss and summarize how COM feedback delay affects different aspects of balance control, including co-contraction and joint stability, as well as balance strategy.

## Materials and Methods

### Musculoskeletal Model and Perturbation Modeling

A generic 2D MSK model was adapted from the gait10dof18musc model available in OpenSim [40] and used in this study, as shown in Figure 1. The model (weight 74.5 kg, height 1.8 m) consists of 9 degrees of freedom (DOFs) and 18 Hill type muscles all with actuation on the sagittal plane [41]. A moving platform weighing 10 kg was added to the model and connected to the ground through a sliding joint allowing the platform to translate only in the anterior-posterior (AP) direction. This sliding joint was controlled to generate a continuous sinusoidal movement, which was set to magnitudes from 0 mm to 80 mm at a frequency of 1 cycle per second (i.e., 1 Hz). The contacts between the feet and the platform were modeled as Hunt-Crossley contact forces [42]. Contact spheres were placed on the toes and heels of both feet, and the surface of the platform was covered by a contact half space. The parameters used to model the contact forces are listed in Table 1.

**Figure 1:**
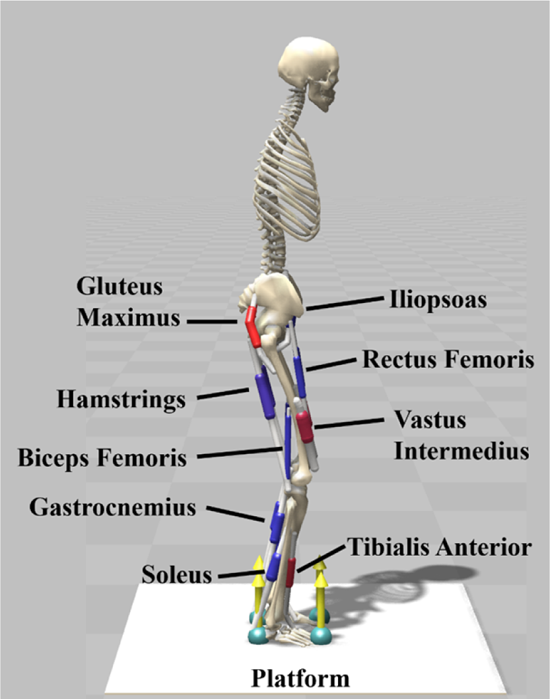
The musculoskeletal model used in the simulations. The nine muscles included on each leg are labeled. The platform moves in a cyclic pattern in the anterior-posterior direction with different amplitudes set by a controller.

**Table 1:** The contact parameters for the model contact forces, based on Hunt-Crossley contact forces (42).

| Parameter | Value |
| --- | --- |
| Stiffness | $2 \times 10^6 \text{ N/m}^2$ |
| Dissipation | 1 |
| Static Friction Coefficient | 0.9 |
| Dynamic Friction Coefficient | 0.6 |
| Viscous Friction Coefficient | 0.6 |
| Transition Velocity | 0.15 m/s |

### Muscle Reflex and COM Feedback Control

To maintain balance under the continuous perturbation, the MSK model was controlled by a three-way feedback control approach as shown in Figure 2. The activations for the model’s muscles were calculated through three different feedback controllers: a low-level muscle length feedback controller, a muscle force feedback controller, and a COM feedback controller. Employing multiple controllers to balance the model reflects the redundancies found in the human body.

**Figure 2:**
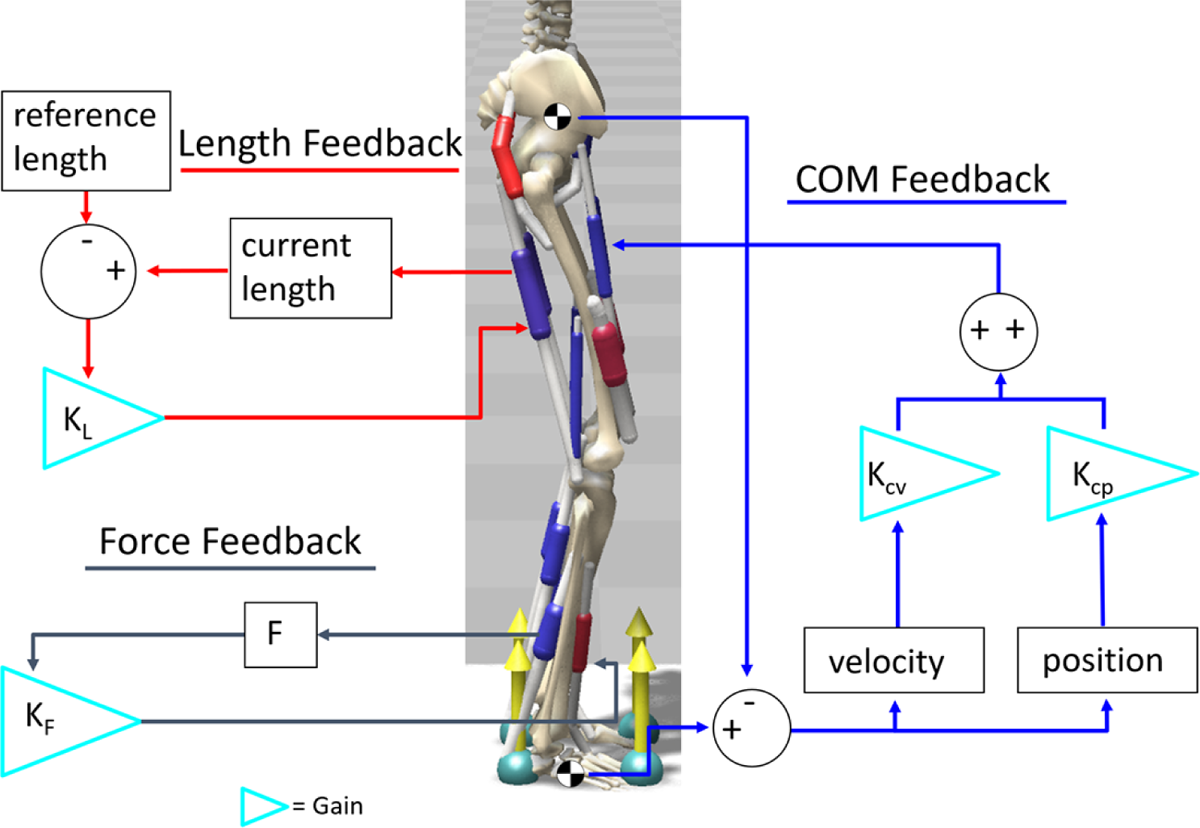
The three-way feedback control approach used to balance the model. Length reflex and force feedback mimic sensory feedback in the muscles, while the COM feedback reacts to the position and velocity of the body and foot COM.

The low-level muscle length feedback controller activates muscles based on changes in muscle fiber length. The muscle fiber length thresholds and activation delays are taken from previous studies [34, 43–45]. The guiding principle behind the controller is that once a muscle is stretched beyond a certain point, reflexes will activate the muscle to keep the body in balance; this is modeled after the muscle spindles [34]. The controller calculates activations using the following equation:

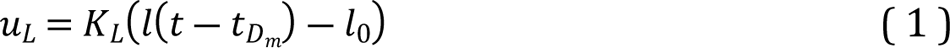

where *u*_*L*_ is the length reflex caused muscle excitation, *l* is the current normalized muscle fiber length, *l*_0_ is length feedback offset, *K*_*L*_ is the length gain, *t* is the current time, and *t*_*Dm*_ is individual muscle reflex time delay (*m* is the muscle index) which ranges from 35 ms to 100 ms [43–45].

The muscle force feedback controller was used to mimic co-contraction, which is a stabilizing factor in the human body. The controller works by detecting the force output of a muscle and then activating the corresponding antagonist muscle to balance the force production on either side of the limb. In simulations, this controller calculates the muscle activation output (*u*_*F*_) using the following equation:

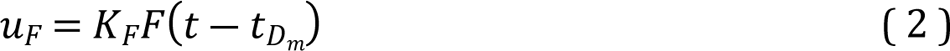

where *F* is the force of an antagonist muscle, *K*_*F*_is the force gain, and *t*_*Dm*_ is the same time delay as used in equation (1). To facilitate the muscle force feedback mechanism, several agonist and antagonist muscle pairs are established and listed in Table 2.

**Table 2:** Co-contractor pairs used in the force feedback controller and co-contraction index calculations.

| Agonist | Antagonist |
| --- | --- |
| Gluteus Maximus and Hamstrings | Iliopsoas |
| Hamstrings and Biceps Femoris | Rectus Femoris and Vastus Intermedius |
| Gastrocnemius and Soleus | Tibialis Anterior |

The COM feedback controller activates muscles based on the model’s whole body COM position and velocity relative to the foot COM position and velocity in the sagittal plane. The purpose of this controller is to keep the model’s COM over its BoS, acting as the vestibular balance component of human balance. The muscle activations determined by the COM reflex controller is defined as follows:

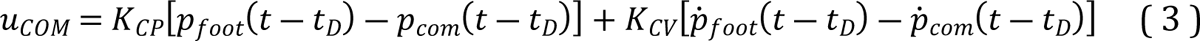

where *p* indicates the position and *p* indicates the velocity of the foot or whole-body COM; and *K*_*CP*_ and *K*_*CV*_are the feedback control gains for position and velocity respectively, and *t*_*D*_is the COM feedback delay. The range of delays used were adapted from the study by McKay *et al*. [46], which looked at sensory feedback and balance in individuals with Parkinson’s disease compared to elderly individuals without Parkinson’s disease; in this study they provided a sensorimotor feedback model that successfully reproduced muscle activations as seen in experimental trials with healthy, aging, and Parkinson’s disease subject groups. Equation (3) differs from the equation used in McKay *et al.*’s study by using a comparison of whole body COM position to foot COM position. The relative position of these two elements informs the stability of the body and in which direction it needs to be corrected. For example, if the body COM moves forward beyond the foot COM, body kinematics need to be adjusted to shift the body COM backwards.

### Optimization and Simulation Setup

The model’s initial state of standing was set to a slightly crouched posture in anticipation of the expected perturbation and better stability: a pelvis tilt forward of 10 degrees, both hips flexed at an angle of 20 degrees, both knees flexed at an angle of 20 degrees and both ankles dorsiflexed at an angle of 10 degrees. Additionally, all initial joint velocities were set to zero at the start of the simulation.

Three fixed COM feedback delay conditions were used to investigate the effect of delayed feedback: *t*_*D*_ = 100, 150, and 200 ms. These delay conditions were modeled to reflect the sensory feedback delay observed in the elderly population [46]. The magnitude of the sinusoidal perturbation applied to the platform was increased from 0 to 80 mm, where the 0 to 10 mm range was incremented by 2 mm and the 10 mm to 80 mm was incremented by 20 mm. A perturbation magnitude of 80 mm was used as the maximum in this series of simulations because magnitudes greater than 80 mm were observed to cause the model to fall. In total, the perturbation magnitudes applied to the model were: 0 mm, 2 mm, 4 mm, 6 mm, 8 mm, 10 mm, 20 mm, 40 mm, 60 mm, and 80 mm. These magnitudes were then repeated for each COM feedback delay condition, resulting in a total of 30 simulations. The frequency of the perturbation was kept at 1 Hz for all simulations. In Figure 3, a graphical representation of the perturbation cycle is shown to illustrate the magnitude and velocity of the perturbation and its four different phases.

**Figure 3:**
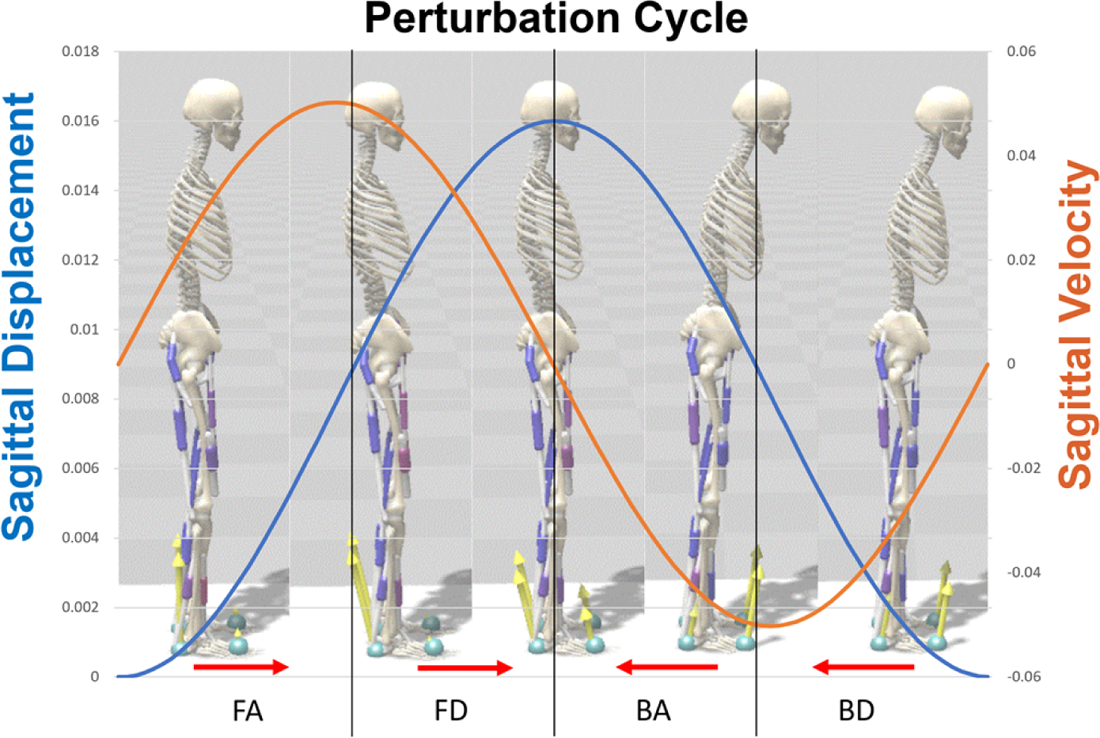
A graphical representation of a single perturbation cycle (80mm magnitude of sinusoidal perturbation with a total 160mm movement in a cycle). Four different phases of the perturbation include FA: forward accelerating; FD: forward decelerating; BA: backward accelerating; BD: backward decelerating.

For each simulation, an optimization was performed to find optimal parameters for *K*_*F*_, *K*_*L*_, *K*_*CP*_, *K*_*CV*_, *l*_0_ such that a defined objective (cost) function is minimized. The objective function seeks to minimize effort (squared sum of muscle activations) while penalizing falling (with termination when the COM height reaches below 0.9 times the initial COM height), large knee angles (>30°), pelvic tilt (>±15°) and pelvic anterior-posterior (AP) translation (>±0.2 m) on the sagittal plane. The objective function to be minimized was defined as follows:

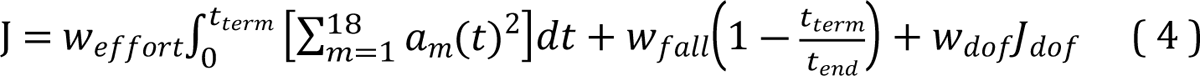

where w is weight (*W*_*fall*_ = 100, *W*_*effort*_ = *W*_*dof*_ = 1), *t*_*end*_ is the maximum simulation duration (set to 10 s), *t*_*term*_ is the simulation termination time when the COM height of the body falls below 0.9 times its initial COM height (*t*_*t*er*m*_ = *t*_*end*_ if the model never falls). *J*_*dof*_is the penalty associated with exceeding preset joint DOF ranges that indicate tendance of balance loss:

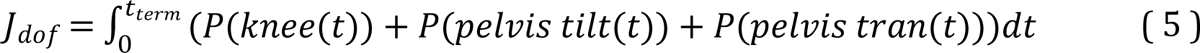

with *P*(*knee*(*t*)), *P*(*pelvis tilt*(*t*)), *P*(*pelvis trans*(*t*)) set to 10 if the knee flexion angle is greater than 30, pelvis title angle is greater than ±15°, and pelvis translation is greater than ±20cm, respectively, and set to 0 otherwise.

All optimizations were run with the commercial Hyfydy version of SCONE [47] for faster computations [48]. The SCONE software package uses Covariance Matrix Adaptation Evolution Strategy (CMA-ES) to optimize model parameters. In principle, CMA-ES functions similar to the survival of the fittest concept in biology, where generations are iterations, genetic traits are parameters, and survival is based on the objective function value [49]. Iterations with more favorable parameters survive to the next generation until an optimal set of parameters is found. For efficiency, the optimal results of one perturbation magnitude were used as the initial guess for the optimization of the next higher perturbation magnitude. For each simulation case, the optimization was run in parallel 60 times with different random seeds and the optimization result with the lowest objective value was chosen as the final optimal result. All optimizations were run on a desktop computer with Intel Xeon 3.7 GHz CPUs and optimization for each case typically converged within an hour.

### Data Processing

The joint angle and muscle activation data was processed by first segmenting the data into one-second cycles, then determining the mean and standard deviation of the data across all the cycles. The first two seconds of the simulation were removed from the analyses, as the muscle activations at the beginning of the simulation were more volatile than those when the model has settled into a more stable postural pattern. This also renders the initial posture less influential on the results after the first two seconds.

The co-contraction index (CCI) of each agonist-antagonist muscle pair (as defined in Table 2) was calculated using the equation from Rudolph *et al*. [50]:

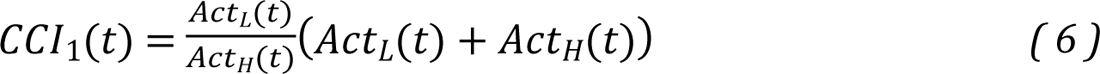

where *t* is time, *ACt*_*L*_ is the lower activation and *ACt*_*H*_ is the higher activation of the muscle pair. This equation gives the time history of CCI during the cycles. To better compare the CCI between different simulations, we integrated the CCI to give a final scalar number conveying the strength of the co-contraction between the muscles for each condition. The CCI can be related to joint stability, with a higher CCI correlating to a less joint stability [50–53].

The balance strategy used by the model in each simulation was determined by calculating the Pearson’s Correlation coefficient for adjacent joint moments (e.g., hip to knee or knee to ankle), using the following equation:

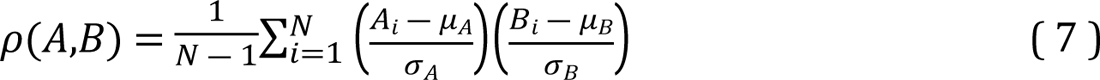

where A and B are the adjacent joint moments, *N* is the number of simulation steps, and μ and σ are the mean and standard deviation respectively. For example, if the ankle and the knee express moments in the same direction, they are in phase and have a positive correlation. If their moments are in opposite directions, they are anti-phase and have negative correlation. The same would apply to the relationship between the hip and knee. Once the correlation coefficients were calculated, the strategy was determined using criteria developed by Yeadon and Trewartha [54] and further expanded upon by Blenkinsop, *et al*. [31], see Table 3 for more information. There are other, more complex ways to determine balance strategy. For example, Taleshi *et al*. also used vector coding to determine strategy, which used relationships of adjacent joint moments, but also added in COM and COP motion [55]. In the end, we chose to use the current approach because of its simplicity and straightforward interpretation.

**Table 3:**
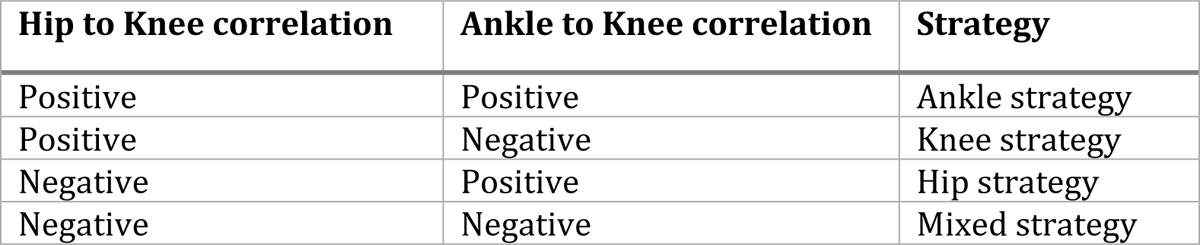
Criteria for determining balance strategy.

| Hip to Knee correlation | Ankle to Knee correlation | Strategy |
| --- | --- | --- |
| Positive | Positive | Ankle strategy |
| Positive | Negative | Knee strategy |
| Negative | Positive | Hip strategy |
| Negative | Negative | Mixed strategy |

## Results

### Joint Angles

Figure 4 shows the joint angles across all large magnitudes of perturbation (10 mm to 80 mm) and all COM feedback delay conditions. The perturbation magnitudes of more than 10 mm resulted in larger ranges of joint angles compared to those below 10 mm; however, interestingly, the largest magnitude perturbance did not always produce the largest joint angles. Overall, the 150 ms delay produced higher peak joint angles than the other two delays and the 100 ms delay resulted in the highest standard deviation of joint angles. Additionally, looking at the absolute joint angle ranges – which reflects the maximum and minimum joint angle deviation from a 0° or neutral joint angle – across all perturbations and magnitudes there appears to be a trend of the joint angle range increasing for larger magnitudes of perturbation for the knee and ankle, but decreasing for the pelvis and hip (Figure 5). The decreased pelvis angle indicates more upright posture and less sway of the upper body during balance since the pelvis and torso are fused together in the model.

**Figure 4:**
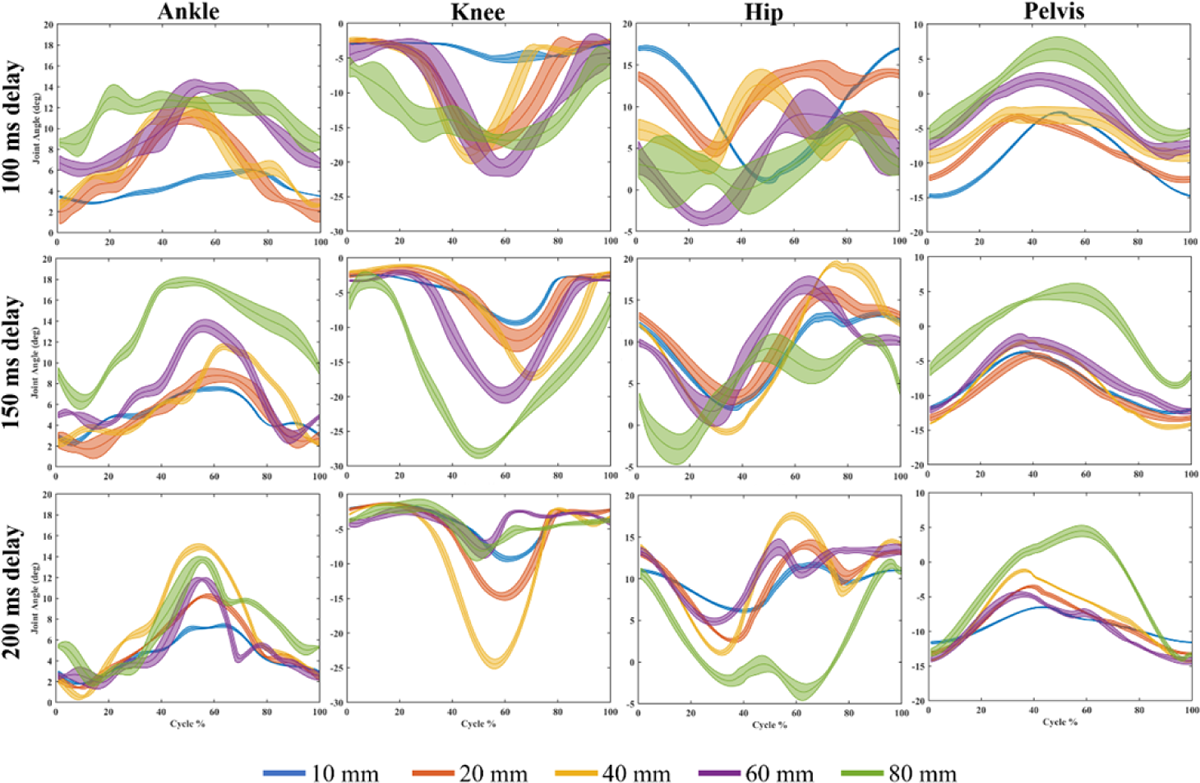
Joint angles for each delay at perturbation magnitudes: 10 mm, 20 mm, 40 mm, 60 mm, and 80 mm. The data is broken down into eight cycles. The solid line is the mean joint angle, and the shaded area represents the mean ± standard deviation region.

**Figure 5:**
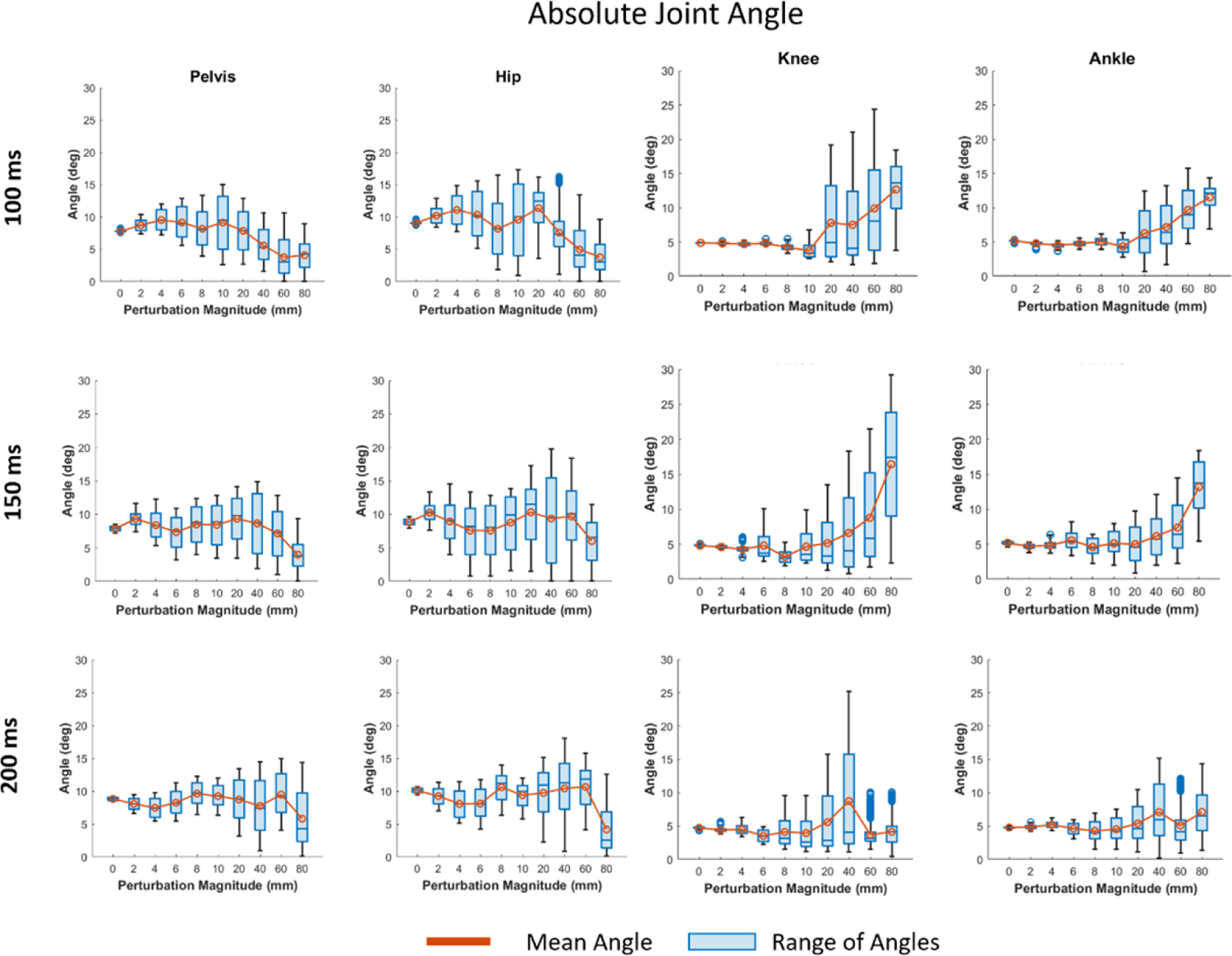
The absolute joint angle range for each joint at each COM feedback delay and perturbation magnitude. The line plot represents the mean joint angle. The blue boxes represent the range of joint angles observed in each simulation.

### Muscle Activations

The COM feedback delay appeared to affect peak muscle activation, as can be seen in Figure 6. This effect is most prominently seen in the anterior muscles (rectus femoris (RF), vastus intermedius (VAS), and tibialis anterior (TA)). Additionally, while the gastrocnemius medialis (GAS) and TA show greater activations in response to longer delays, the soleus (SOL) keeps a similar range of activation and has a more chaotic pattern. While the gluteus maximus (GM) and biceps femoris (BF) muscles were modeled, and included in all simulations, the activations of these muscles were minimal. Both muscles are monoarticular muscles that span the hip and knee joints, respectively. In contrast, biarticular muscles such as RF, GAS, and hamstrings (HM) spanning the hip and knee joints have relatively high activation.

**Figure 6:**
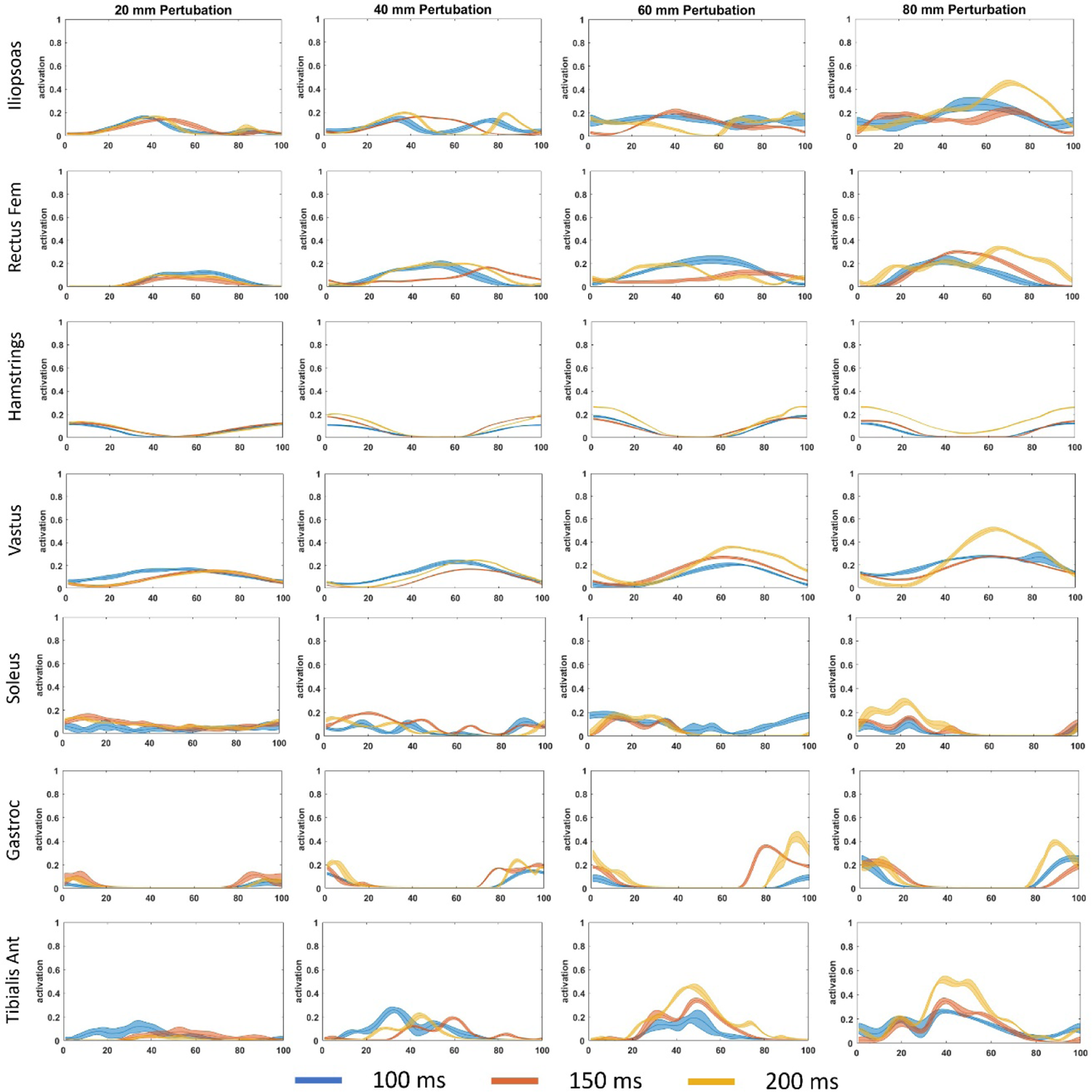
Normalized muscle activation (mean and standard deviation) over the averaged perturbation cycle for each delay (100 ms, 150 ms, 200 ms) and magnitude (20 mm to 80 mm). Gluteus Maximus and Biceps Femoris are not included because their activations were very low.

To further explore muscle activations, we calculated the CCI (Eq. (4)) for antagonist/agonist pairs crossing the knee, hip, and ankle joints (Table 2). Once again, because the BF and the GM activations were very low, they were not included in the presented data in Figure 7. While the CCIs for lower magnitude perturbations (less than 80 mm) show less significant differences across the three delay conditions, the 80 mm delay condition produced higher CCI, especially for the 200ms delay condition.

**Figure 7:**
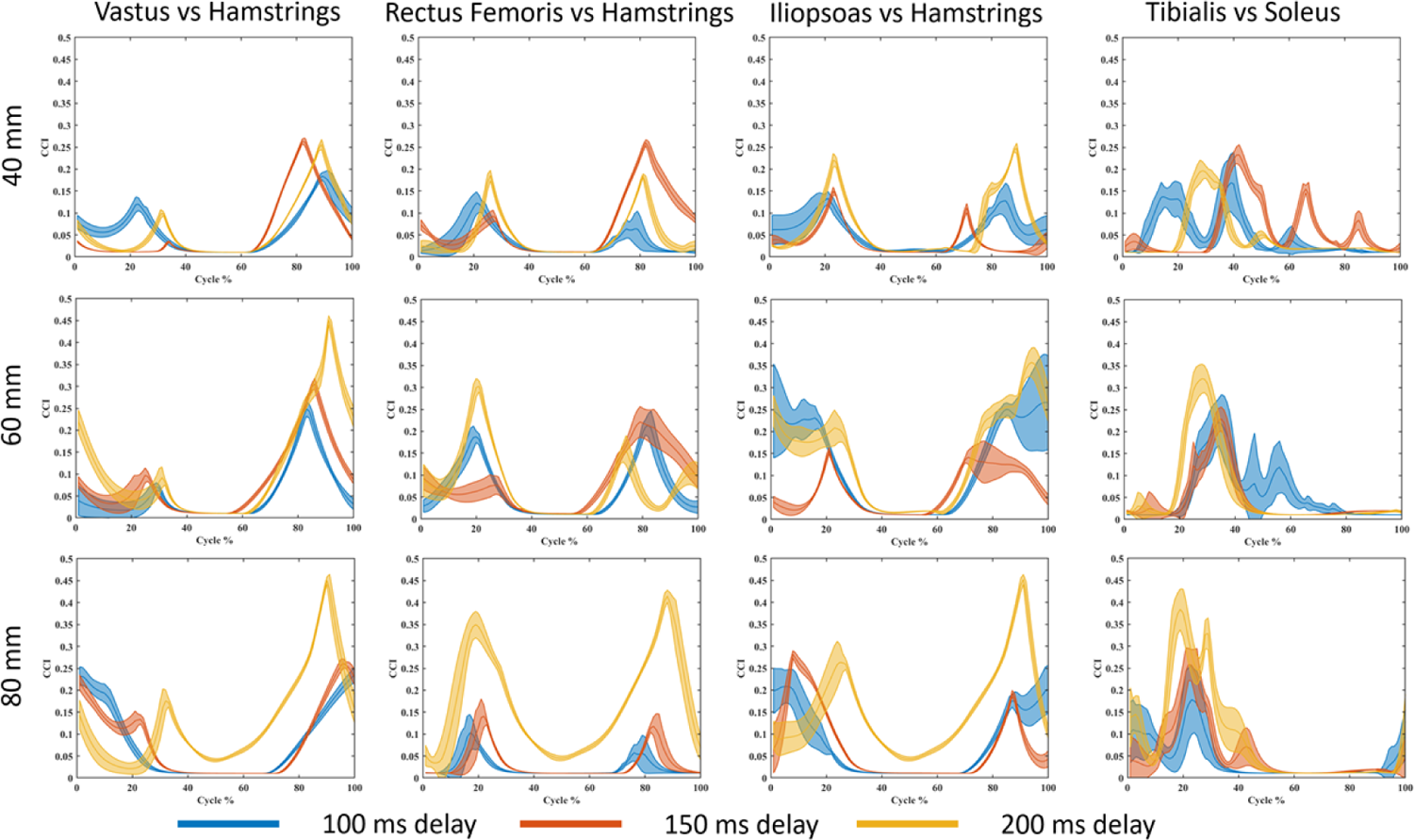
Co-contraction index calculated based on the Rudolph equation for perturbation magnitudes of 40 mm to 80 mm.

A more intuitive way to compare CCI values and observe differences across conditions is to integrate the CCI curve and compare the resulting areas rather than discrete time series [50, 52]. The results from integrating the CCI are presented in Table 4; the data are grouped by COM feedback delay conditions for each co-contraction pair at each perturbation magnitude. In Table 4, it can be observed that the CCI is in general much higher at 60mm and 80 mm perturbations for the 200 ms delay, whereas the pattern is less clear at perturbations of lower magnitude.

**Table 4:** Co-contraction index (CCI) integrals for muscle pairs with major activations. Vastus Intermedius (VAS), Hamstrings (HM), Rectus Femoris (RF), Iliopsoas (IL), Tibialis Anterior (TA), Gastrocnemius Medial (GAS), and Soleus (SOL). Darker shades of green represent higher CCI values and therefore less joint stability.

|  |  | 0 mm | 2 mm | 4 mm | 6 mm | 8 mm | 10 mm | 20 mm | 40 mm | 60 mm | 80 mm |
| --- | --- | --- | --- | --- | --- | --- | --- | --- | --- | --- | --- |
| 100 ms | VS - HM | 2.34 | 4.33 | 5.96 | 5.65 | 6.17 | 7.10 | 7.49 | 6.34 | 5.42 | 7.90 |
|  | RF-HM | 1.60 | 1.49 | 1.47 | 1.33 | 1.33 | 1.33 | 2.82 | 3.34 | 6.73 | 2.38 |
|  | IL-HM | 1.60 | 1.49 | 1.43 | 1.38 | 1.37 | 1.54 | 3.42 | 5.58 | 12.35 | 7.60 |
|  | TA-GM | 2.00 | 1.67 | 1.69 | 1.62 | 1.42 | 1.74 | 1.51 | 1.63 | 1.33 | 4.88 |
|  | TA-SOL | 1.20 | 1.22 | 1.23 | 1.31 | 1.48 | 1.25 | 3.41 | 4.89 | 5.60 | 4.30 |
| 150 ms | VS - HM | 2.76 | 4.10 | 4.02 | 3.88 | 3.34 | 6.06 | 6.59 | 5.83 | 8.29 | 8.36 |
|  | RF-HM | 1.61 | 1.41 | 2.21 | 4.68 | 2.97 | 3.58 | 2.91 | 7.21 | 8.37 | 3.19 |
|  | IL-HM | 1.61 | 1.39 | 1.42 | 2.01 | 2.39 | 4.34 | 4.38 | 3.47 | 6.42 | 7.30 |
|  | TA-GM | 1.79 | 1.46 | 1.60 | 1.75 | 1.67 | 1.71 | 1.62 | 2.14 | 1.60 | 3.50 |
|  | TA-SOL | 1.18 | 1.17 | 1.23 | 1.17 | 1.25 | 1.83 | 3.54 | 6.11 | 4.26 | 5.37 |
| 200 ms | VS - HM | 5.57 | 5.39 | 6.97 | 7.98 | 8.78 | 5.83 | 5.84 | 6.25 | 11.36 | 12.83 |
|  | RF-HM | 3.02 | 4.07 | 4.68 | 6.75 | 6.73 | 2.87 | 2.88 | 4.67 | 7.94 | 16.79 |
|  | IL-HM | 1.33 | 3.88 | 4.29 | 6.22 | 4.73 | 3.44 | 4.47 | 7.10 | 14.51 | 15.64 |
|  | TA-GM | 1.67 | 1.64 | 1.57 | 1.87 | 1.76 | 1.56 | 1.56 | 1.31 | 1.83 | 4.75 |
|  | TA-SOL | 1.18 | 1.60 | 1.23 | 1.15 | 1.69 | 2.46 | 2.00 | 4.36 | 5.86 | 8.91 |

### COM and COP

A measure of a person’s stability can be determined by subtracting the COM position from COP position [56]. A difference between the COP and COM close to zero would indicate higher stability, whereas a difference further from zero would indicate less stability. Figure 8 shows that the COP in the AP direction had a wider range than the COM, and that range increased with both larger magnitude perturbations and longer COM feedback delays. Figure 9 shows the mean stability, calculated from the difference between COP and COM, for the larger magnitudes of perturbation (>10 mm) at each of the feedback delay conditions. Overall, the greatest peaks of instability (furthest values from zero) are seen in the larger perturbation conditions; however, for the 100 ms delay, there is a less prominent difference among the perturbance magnitudes.

**Figure 8:**
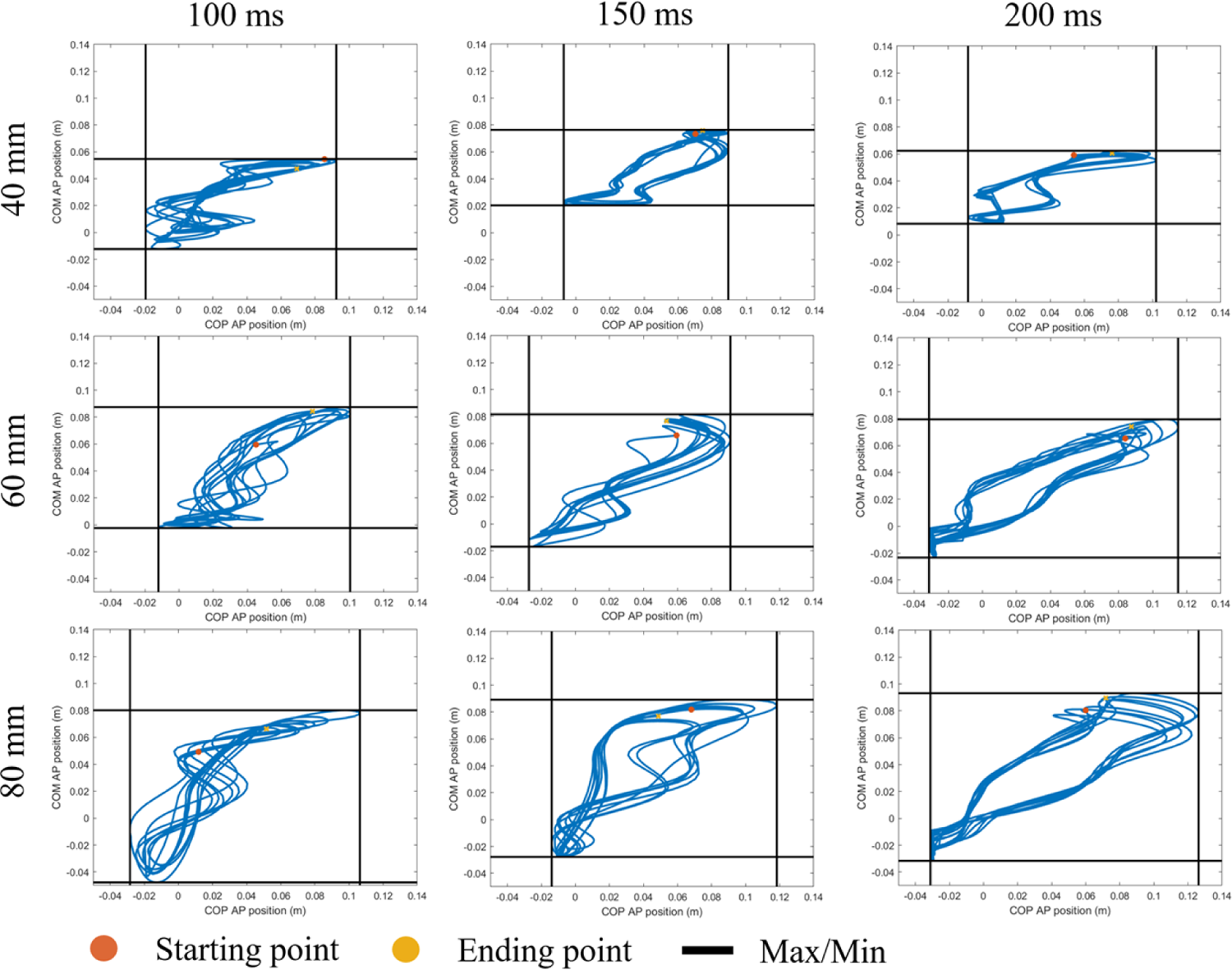
The COP vs COM for each delay in the anterior-posterior (AP) direction for the highest three perturbations (40 mm, 60 mm, and 80 mm). The black solid lines represent the maximum and minimum position of COM and COP. The X-axis represents COP values while the Y-axis is the COM values.

**Figure 9:**
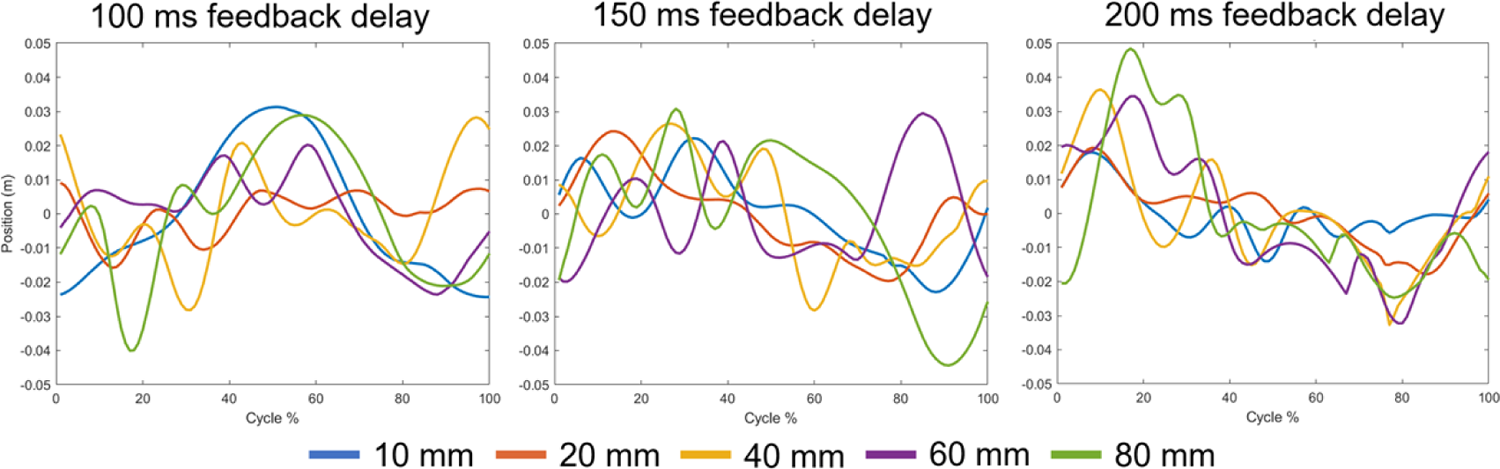
The mean difference between COP and COM AP position for each delay and the perturbance magnitudes greater than 10 mm.

### Balance Strategy

Joint moments across the different perturbation magnitudes showed overall increasing joint moments with increasing magnitudes (Figure 10). The joint moment patterns do appear to change for different COM feedback delays, with an overall higher instance of ankle strategy being used with longer COM feedback delays. Balance strategy during the perturbation cycles was calculated by comparing Pearson’s correlation in the moments of adjacent joints and the criteria shown in Table 3. The results for balance strategy during the perturbation cycles are shown in Figure 11 for the four highest perturbations and all three delays. As can be observed, prolonged COM feedback delays altered the balance strategies during different phases of the perturbation cycles. While the 150 ms delay causes more frequent use of hip strategy during the forward platform movement phase with high amplitudes, the 200 ms delay appears to favor the ankle strategy more. Table 5 summarizes the dominate balance strategy used for each condition determined by the amount of time the strategy is used. Table 6 shows the overall most used balance strategy and the percentage of time spent using each balance strategy. The amount of time was computed by determining the balance strategy – as discussed in the methods – for each time step, then calculating the percentage of each strategy used.

**Figure 10:**
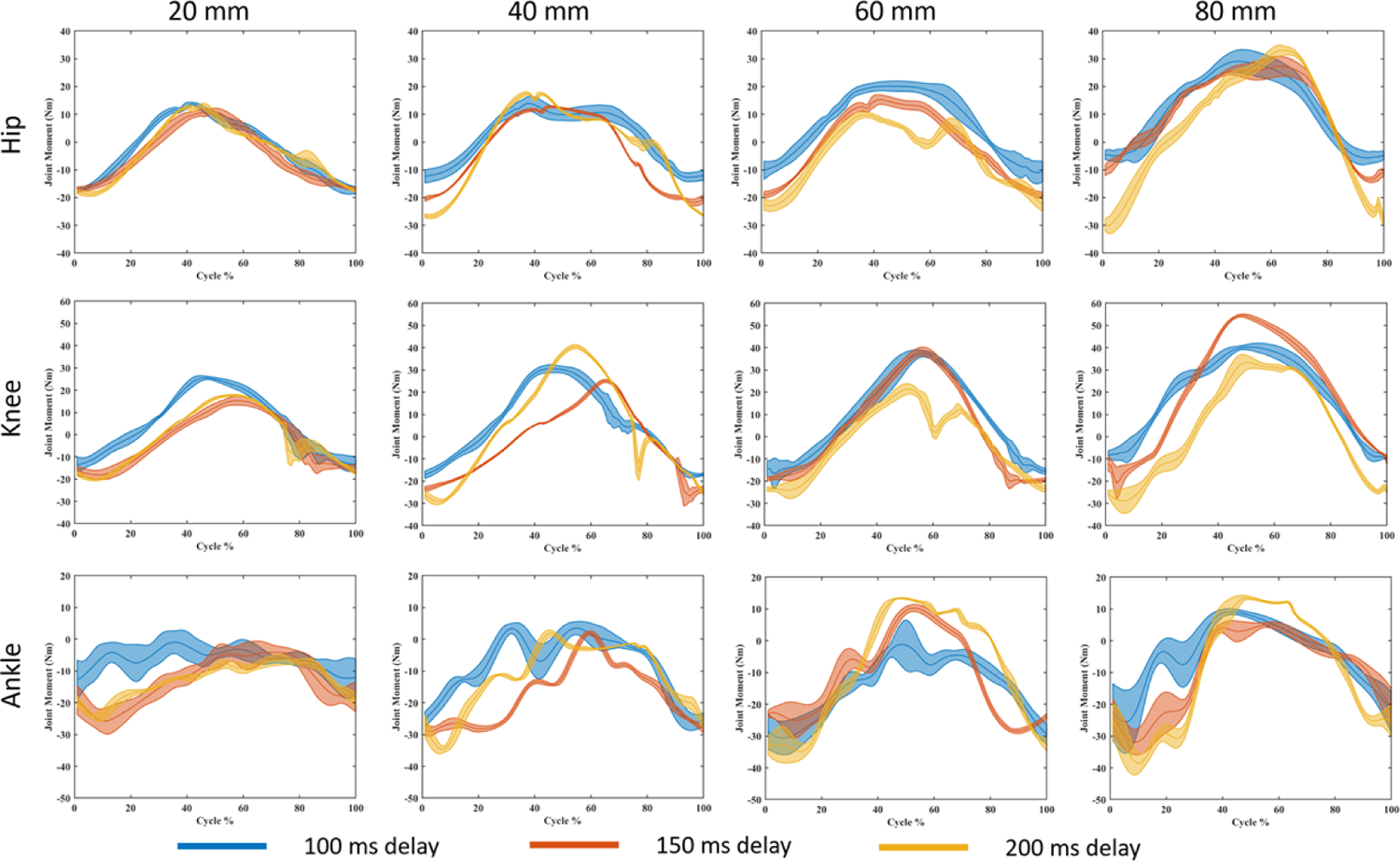
Joint moments for the hip, ankle, and knee at the largest four perturbation magnitudes. The line represents the mean, and the shaded area is the standard deviation.

**Figure 11:**
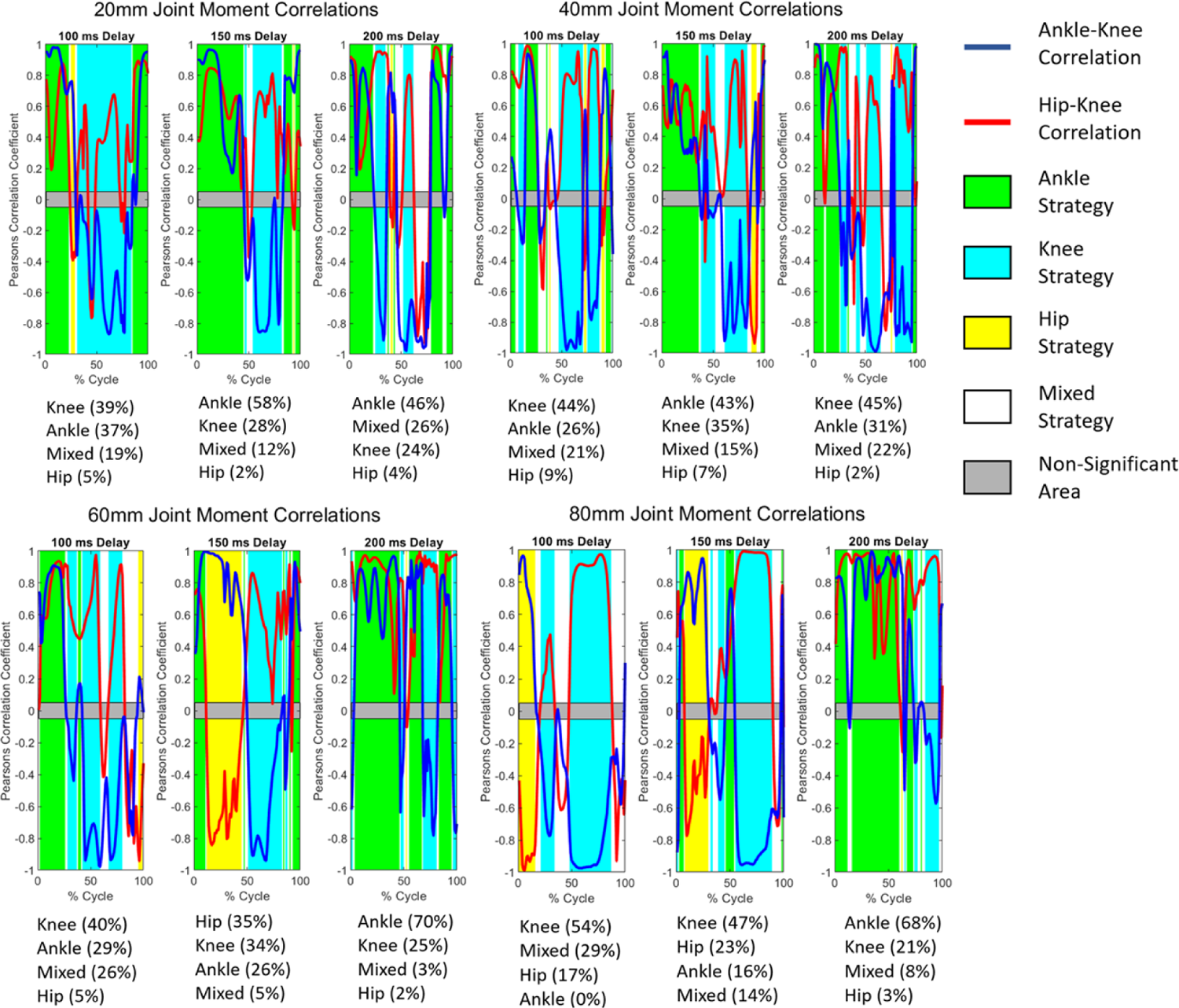
Mean balance strategies employed during different phases of perturbation cycles under 4 large perturbation amplitudes (20,40,60, and 80mm) and three COM feedback delay conditions. The balance strategy was determined at each time step of the segmented data based on a comparison of Pearson’s correlation of adjacent joint moments. See Table 3 for criteria for balance strategy. A Pearson’s correlation coefficient between 0.05 and −0.05 was considered non-significant.

**Table 5:** Dominate balance strategy for each perturbation magnitude (in mm) at each feedback delay (in ms).

|  | 0 | 2 | 4 | 6 | 8 | 10 | 20 | 40 | 60 | 80 |
| --- | --- | --- | --- | --- | --- | --- | --- | --- | --- | --- |
| 100 | Ankle | Ankle | Ankle | Ankle | Ankle | Ankle | Knee | Knee | Knee | Knee |
| 150 | Ankle | Ankle | Ankle | Hip | Ankle | Ankle | Ankle | Ankle | Hip | Knee |
| 200 | Ankle | Ankle | Ankle/Hip* | Ankle | Ankle | Ankle | Ankle | Knee | Ankle | Ankle |

\* indicates equal percentage use of two or more strategies

**Table 6:**
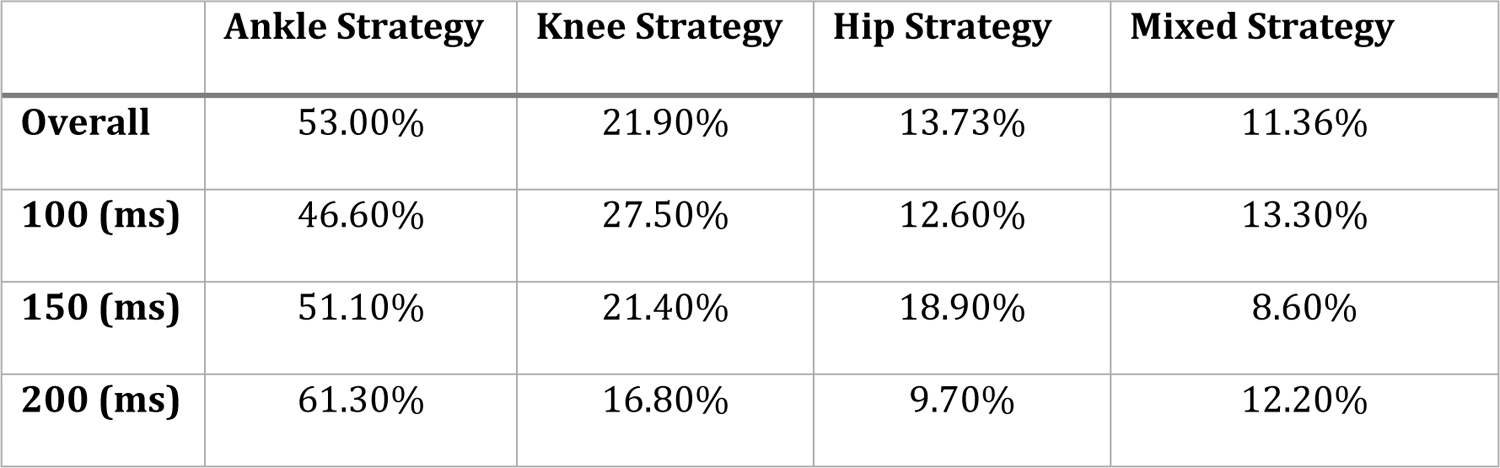
Overall strategy used across all conditions and three delays, computed by calculating the balance strategy for each time step and then finding the percentage of time spent in each strategy.

## Discussion

In this study, the effects of COM feedback delay were evaluated using a MSK model. Continuous, sinusoidal perturbations at ten magnitudes (0-80 mm) were applied to the model via the sliding platform on which it stood. Further, the simulations were repeated for three COM feedback delays: 100 ms (fast/healthy), 150 ms (slow/aging/slightly delayed), and 200 ms (slowest/severely delayed). Analysis of the data suggests that the differences in joint angles, muscle activations, and COP and COM positions for each level of delay indicate a multifaceted effect of COM feedback delay on the balance mechanisms.

### Muscle Activations

In Figure 6, the peak muscle activation happens during the backward deaccelerating phase in some muscles – most notably in the RF, which is an important stabilizing muscle. The delay in COM feedback means that the body cannot respond to perturbations at the near-instantaneous rate seen in younger populations, and therefore proactive strategies for balance control are important. For example, anticipating and preparing for a perturbation, by bracing or stepping, can help reduce the risk of falls. Additionally, Figure 6 showed muscle activations were larger overall at longer feedback delay conditions. In a study by Chung *et al*. [20] on neuromuscular age-related changes, the authors found that for the same amount of activation, elderly subjects produced lower joint torques than their younger counterparts. To address these problems, training programs that focus on improving balance reactions and reducing the muscle reflex delay can be beneficial for fall prevention in older adults and individuals with balance impairments. In a study by Duncan *et al*. [57], the authors show that a person’s past experience with instability can affect how quickly they adapt to new perturbances, which supports the idea that training can help individuals adapt to perturbances more quickly.

CCI calculations give insight into the amount of joint stabilization being produced by agonist/antagonist pairs of muscles surrounding a joint [50, 52]. One method to calculate the CCI is by using muscle activations (via EMG) or joint moments. The two commonly used equations for calculating CCI are the Falconer and Winter [52] equation and the Rudolph [50] equation. In this study we chose to use the Rudolph equation based on a 2021 study by Li *et al*. [51], which found that the Rudolph equation produces more accurate results, especially when using muscle activations as inputs – which was the case in this study. The Rudolph CCI equation (Eq. (6)) produces a time curve that gives the CCI at any given time in the cycle. That curve can then be integrated to give the area, which gives a more straight-forward comparison of the result (Table 4). The CCI calculated for this study showed higher CCI at 80 mm of perturbance (Figure 7), which is logical given that it is the largest magnitude perturbation at which the model could still balance. Additionally, at 80 mm, the CCI for 200 ms feedback delay was higher than the other delay conditions, showing higher CCI than seen in the same muscle pairs during walking studies [58]. According to Rudolph *et al*. [50], higher CCI corresponds to lower joint stability. In this study, the tendency of higher CCI seen in longer COM feedback delays suggests that the inherent longer sensory feedback delay seen in the elderly population may be contributing to low joint stability and balance difficulties. Joint stability can be increased by a variety of methods, including neural feedback control which can be affected by delay, impedance control by relying on the passive properties of muscles, and co-contraction of antagonist/agonist muscle pairs which is effective but metabolically inefficient [59]. On the contrary, studies have also found lower joint stability is not an inherent factor of aging [8, 60, 61]. This calls for further research into the relation between sensory feedback delay and joint stability.

### Balance Strategies

The human body uses the hip, knee, and ankle joints to shift the COP to control the COM position. The moments and angles of these joints and how they relate to each other is how researchers determine balance strategy. The criteria used to determine balance strategy in this study are laid out in Table 3. The balance strategy used is sensitive to the magnitude of perturbation. To maintain balance, ankle strategy is used for fine adjustments, in response to mild perturbations, whereas more severe perturbations require rough adjustments from knee or hip strategy [32, 33, 38]. This pattern is reflected in our simulation results, where we saw ankle strategy was dominant for the majority (94%) of conditions below 20 mm across all delay conditions (Table 5) and was used 53% of the time over all simulations (Table 6). In the 2017 study by Blenkinsop *et al*. [31], the authors investigated balance strategies used to adapt to discrete perturbations and saw an ankle strategy being used for the majority of the time and a hip strategy being used when perturbations were too large for the ankle. Likewise, Taleshi *et al*. [55] found that an ankle strategy is used in lower frequency perturbations, however they found a knee strategy being used in higher frequency perturbations instead of a hip strategy. The Blenkinsop study used discrete perturbations, whereas the Taleshi study used continuous perturbations, which might explain the differences in their findings. In our study, we used continuous perturbations and while we found ankle strategy to be dominant, knee strategy was used more than hip strategy.

Table 6 reveals an interesting trend: the use of ankle strategy increases with longer feedback delays. In a 2016 balance simulation study by Versteeg *et al.* [62], it was noted that while hip strategy predominated when minimizing the center of mass (COM) excursion, ankle strategy was preferred for minimizing trunk angle. Our model, with its locked lumbar joint, essentially equates pelvis rotation with trunk angle. We observed a modest increase in average trunk angle with lengthier feedback delays (7.38°, 7.89°, and 8.33° for increasing delays, respectively). This might explain the heightened ankle strategy use in longer delays, reflecting Versteeg et al.’s findings. In a 2011 study by Gurses *et al*. [63], the authors compared balance strategies of young adults vs elderly for different frequencies of perturbation and found that young adults predominantly used hip strategy, whereas elderly individuals employed a mix of hip and ankle strategies used by elderly. This implies an age-related shift towards ankle strategy in response to perturbations. In another study by Taleshi *et al.* [55], it was found that ankle strategy dominated when perturbance frequency was below 2.11 Hz. Our study’s low perturbation frequency, along with increased COM feedback delay (indicative of aging), likely contributed to the increased use of ankle strategy.

The type of balance strategy being used appears to vary with phases of the perturbation cycle, as can be seen in Figure 11. For the majority of the simulations in Figure 11, the ankle strategy is predominately used during the early phase of the perturbation cycle when the platform is moving forward, while the knee strategy is mostly used during phases characterized by platform backward movement. Notably, existing research on continuous perturbation lacks a comprehensive categorization of balance strategies according to perturbation cycle phases, necessitating further investigation. Subsequent studies, particularly those involving experimental validation, are imperative to elucidate the extent to which perturbation direction influences balance strategy. Factors such as speed and the anticipation of speed transitions should also be explored to comprehensively understand the interplay of these elements in shaping adaptive postural responses.

### COM and COP

It has been established in previous studies, that for the human body to maintain upright balance, the COM should be kept within the BoS [64, 65]. In most scenarios, this is accomplished by the body shifting the COM back to a more central location related to the COP, similar to what is seen in the inverted pendulum on a cart design. Because of this relationship, one viable way to evaluate balance is to compare the COM location to the COP location, as we did in Figure 8. It can be seen in those graphs that the COP has a wider range than the COM – which is what is expected in simulations where the model was able to maintain its balance because the COP is able to shift beyond the COM to move it back within the BoS. It can also be seen that the ranges of the COM and COP positions are larger as the magnitude perturbation gets larger, suggesting less stability and a greater chance of loss of balance. The trend of a larger COP range at larger perturbations was also seen in a study by Camernik *et al*. [39], in which participants were subjected to anteroposterior perturbations with and without holding on to a supportive handle.

### Limitations

While most studies agree that lower limbs play a heavy role in balance recovery, the upper limbs do contribute as well. Studies on upper limb amputees have shown that an upper limb loss puts people at a higher risk for falls, with or without prothesis use [66]. Additionally, pronation and supination in the feet have been correlated with poor postural stability [67, 68]. These findings suggest that a model with more joints than the ankle, knee, and hip may give a more complete picture of the balance strategies of an individual.

MSK simulations can be limited by the controllers used to mimic the CNS. Researchers are constantly developing new ways to express mathematically how the human body controls balance. In this study, we used multiple controllers simultaneously to reflect the redundancy of the CNS. Other researchers claim that more complex controllers that more precisely mimic the nervous system should be used to obtain more true-to-life results [69]. Still others suggest that a more accurate portrayal of human balance can be found through momentum-based control, rather than COM feedback [70]. Continuing research is needed to determine the best strategy for modeling human balance and obtaining the most accurate to real life results.

Our optimization approach might not always yield the true global optimum, and outcomes can vary even under identical optimization scenarios. While the CMAES algorithm we employ is a global optimization technique, it is not immune to occasionally converging on local minima. To mitigate this, we conducted 60 parallel optimizations for each of the 30 simulation cases and selected the result with the lowest objective function value. Intriguingly, we observed that optimal solutions with similar objective function values could sometimes lead to distinct movement patterns and balance strategies. This observation prompts a critical question about the selection of the most realistic solution. To enhance the reliability of our results and reduce their sensitivity on a single solution, we could potentially adopt an approach similar to how experimental data from multiple subjects are handled. This would involve using a set of optimal solutions and reporting their mean and standard deviation, thereby providing a more comprehensive view of the possible solutions.

## Conclusions

In summary, our study introduced a novel three-way muscle control approach for balancing and investigated how COM sensory feedback delay affects elderly’s balance capability and strategies. The key findings of this research include: 1) Longer COM feedback delay resulted in increased muscles activations and co-contraction, indicating weakened joint stability; 2) Prolonged COM feedback delays led to noticeable shifts in balance strategies during perturbed standing. Under low-amplitude perturbations, the ankle strategy was predominantly used, while higher amplitude disturbances saw more frequent employment of hip and knee strategies. Additionally, prolonged COM delays altered balance strategies across different phases of perturbation, with a noticeable increase in overall ankle strategy usage. These findings underline the adverse effects of prolonged feedback delays on an individual’s stability, necessitating greater muscle co-contraction and balance strategy adjustment to maintain balance under perturbation. Our findings advocate for the implementation of training programs tailored to enhance balance reactions and mitigate muscle feedback delays within clinical or rehabilitation settings, which hold promise for preventing falls in both elderly people and individuals with balance impairments.

## Data Availability

The model and simulation file as well as result data are available at https://github.com/NJITBioDynamics/ModelDataPerturbedBalanceControl/

## Acknowledgements

CDC/NIOSH, Grant contract number: 75D30120P08812.

